# Proteome biology of primary colorectal carcinoma and corresponding liver metastases

**DOI:** 10.1101/2021.10.03.462921

**Authors:** Matthias Fahrner, Peter Bronsert, Stefan Fichtner-Feigl, Andreas Jud, Oliver Schilling

**Author notes:** To whom correspondence should be addressed: Breisacher Straße 55, D-79106 Freiburg (Germany).

## Abstract

Colorectal adenocarcinomas (CRC) are one of the most commonly diagnosed tumors worldwide. Colorectal adenocarcinomas primarily metastasize into the liver and (less often) into the peritoneum. Patients suffering from CRC-liver metastasis (CRC-LM) typically present with a dismal overall survival compared to non-metastasized CRC patients. The metastasis process and metastasis-promoting factors in patients with CRC are under intensive debate. However, CRC studies investigating the proteome biology are lacking. Formalin-fixed paraffin-embedded (FFPE) tissue specimens provide a valuable resource for comprehensive proteomic studies of a broad variety of clinical malignancies. The presented pilot study compares the proteome of primary CRC and patient-matched CRC-LM. The applied protocol allows a reproducible and straightforward identification and quantification of over 2,600 proteins within the dissected tumorous tissue. Subsequent unsupervised clustering reveals distinct proteome biologies of the primary CRC and the corresponding CRC-LM. Statistical analysis yields multiple differentially abundant proteins in either primary CRC or their corresponding liver metastases. A more detailed analysis of dysregulated biological processes suggests an active immune response in the liver metastases, including several proteins of the complement system. Proteins with structural roles, e.g. cytoskeleton organization or cell junction assembly appear to be less prominent in liver metastases as compared to primary CRC. Immunohistochemistry corroborates proteomic high expression levels of metabolic proteins in CRC-LM. We further assessed how the *in vitro* inhibition of two in CRC-LM enriched metabolic proteins affected cell proliferation and chemosensitivity. The presented proteomic investigation in a small clinical cohort promotes a more comprehensive understanding of the distinct proteome biology of primary CRC and their corresponding liver metastases.

## Introduction

Colorectal cancer (CRC) is still one of the most diagnosed cancers worldwide, presenting the third-highest prevalence in men^1,2^. Over 1.1 million new CRC cases were diagnosed in 2020 with over half a million CRC-related deaths worldwide, making CRC the second leading cause of cancer death^2,3^ Unhealthy diet, obesity, smoking, lack of physical activity, and genetic predisposition are established risk factors for CRC^4,5^. Typically, CRC develops over several years, starting as benign adenomatous polyps becoming advanced adenoma with high-grade dysplasia, and then progresses to invasive cancer^6^. Multiple consecutive changes on a genetic level are thought to drive the conversion from normal epithelium to malignant tissue^7^. Early detection and removal of colonic polyps and advances in primary and adjuvant therapy are paramount improvements declining the mortality and increasing patients’ 5-year survival in the past decades^8^. Surgical resection represents the preferred therapeutic method of CRC (stage I to III) providing a potentially curative option. Downstaging with neoadjuvant chemotherapy is a possibility for initially unresectable tumor stages^9,10^ The postoperative outcome depends on the clinical, molecular, and histological features of the disease. The strongest prognostic factor is the pathological stage of the resected tumor. Size, presence of distant metastasis, lymph node positivity, and perineural invasion are substantial^11,12^. CRC can spread lymphatic, hematogenous, contiguous, and in a final stage transperitoneal. The most common metastatic sites are the regional lymph nodes, the liver, and the lungs. Usually, the liver is the first metastatic site because of the venous drainage via the portal vein system. Up to 25% of all patients carry liver metastases at the time of diagnosis, while over 30% develop metastases after resection of primary CRC^4,13^. The 5-year survival rate of patients with metastasizing CRC is decreased by eight times compared to patients with local CRC^13^. Due to the development of refined surgical techniques, such as two-stage hepatectomy, preoperative portal vein embolization, and down-sizing chemotherapy, the number and extent of resections of liver metastasis in CRC increased in the last couple of years steadily and showed a positive result towards long-term survival^9,14^. On a molecular level, CRC is a heterogeneous disease, as the majority of all CRCs occur sporadically (~ 70%), caused by somatic mutations^1,15^. Microsatellite instability, mismatch repair deficiency, APC, RAS, and BRAF mutations are relevant and determined in the standard histopathological examination. In the last couple of years, the growing knowledge of molecular pathogenesis plays an important role to improve targeted treatments^16,17^. Multimodal therapy concepts are well-established with combination chemotherapy and targeted biologic agents. Subsequently, significant improvements in survival were achieved^18,19^.

Formalin-fixation and paraffin-embedding (FFPE) represents the most commonly used preservation method for clinical tissue specimens worldwide^20^. This has led to vast tissue archives, storing tissue specimens of a broad variety of malignancies. Continuous protocol development and optimization rendered FFPE tissue readily accessible for proteomic investigations^21–24^. This has prompted an increasing number of clinical proteomic studies investigating a multitude of malignancies, including rare genetic disorders benefitting from thorough FFPE tissue archives^25–28^. Furthermore, some of the FFPE protocols have been shown to yield comprehensive proteome coverage, reaching over 8.000 identified proteins in single measurements, even with minor amounts of tissue material^21,28,29^.

Many malignant tumors have predominantly been studied on the genetic and transcriptomic level, mainly due to the broad availability and enhanced coverage of those techniques. However, with proteins being the effectors within cells and tissues, representing the products of transcriptomic and translational processes, the importance of a detailed investigation of the proteome becomes evident. Proteomic profiling of CRC and its metastases as the functional translation of the genome is challenging but has a great potential to identify proteins that are linked to tumor progression^30^. In this study, we compare the primary tumor with patient-matched liver metastasis.

We describe the distinct proteome biology of primary CRC and their corresponding liver metastases. Further, this project highlights the practicability and feasibility of the previously published direct trypsinization (DTR) protocol^5^. An activated immune response in the liver metastases is highlighted by the significant upregulation of several components of the complement system. The explorative proteome characterization of primary CRC and liver metastases is complemented by follow-up experiments investigating the spatially distributed expression of significantly dysregulated proteins using immunohistochemistry. Furthermore, an increase in sensitivity using tissue macro dissection for the analysis of primary tumors and their distant metastasis is shown. These characteristics and proteomic profile may lead to uncovering diagnostic and prognostic markers to improve the underlying mechanisms of tumor development and progression.

## Material and Methods

### Ethics statement

The study was approved by the Ethics Committee of the University Medical Center Freiburg (504/17). Patients gave written informed consent before inclusion into the study.

### Patient cohort

Seven patients diagnosed with colorectal carcinoma (CRC) and CRC-LM were included in the study. All patients were operated for CRC and CRC-LM between 2014 and 2016 at the Department of General and Visceral Surgery, University Medical Center Freiburg, Germany. The cohort comprises tissue specimens from five male and two female patients with an age ranging between 48 and 75 years. Further details including tumor localization, tumor grading, and TNM classification are summarized in **Table 1**. Patient data, raw LC-MS/MS data, and analysis result files are available at the European Genome-phenome Archive for appropriate research use (https://ega-archive.org; EGAS00001005641). As patient-centric proteomic data is increasingly regarded as sensitive, personal data^31^, EGA requires adherence to a data access agreement. The data access agreement for this dataset corresponds to the “Harmonised Data Access Agreement (hDAA) for Controlled Access Data” as brought forward by the “European standardization framework for data integration and data-driven in silico models for personalized medicine – EU-STANDS4PM”.

**Table 1.** Clinical Characteristics of the Patient Cohort.

| Patient No. | Localization of primary tumor | Grading | TNM Classification |
| --- | --- | --- | --- |
| 1* | rectosigmoid | ACA (G2) | ypT3 pN0 (0/25) pM1 (HEP) |
| 2 | ascending colon | ACA (G2) | pT3 pN1a (1/18) pM1 (HEP) |
| 3 | rectosigmoid | ACA (G2) | pT3 pN2b (8/31) pM1 (HEP) |
| 4 | sigmoid colon | ACA (G2) | pT4b pN2b (9/21) pM1 (HEP, PER) |
| 5 | sigmoid colon | ACA (G2) | pT4a pN1b (3/13) pM1 (HEP, PER) |
| 6 | ascending colon | ACA (G2) | pT4b pN0 (0/28) pM1 (HEP, PER) |
| 7 | sigmoid colon | ACA (G2) | pT3 pN0 (0/13) pM1 (HEP) |

This study comprised 7 patients with histologically confirmed primary colorectal cancer. All patients had liver metastasis (HEP) and three had additional peritoneal carcinoma (PER) at primary diagnosis. All tumors were moderately differentiated (G2). \*Material after chemotherapy (3 cycles).

### Tissue Collection, Fixation, and tissue macro dissection

Tissue specimens were harvested during surgical removal of primary and metastatic tumors and put in formalin solution immediately after surgical removal. All tissue specimens were gross sectioned, processed, and stained according to routine protocols. For proteomic investigation 10 μm thick tissue slices were automatically deparaffinized and stained for hematoxylin as previously described^21,32^. Macroscopical tumor dissection was performed by an experienced pathologist. Finally, the tumorous tissue of each sample was transferred into a fresh 1.5 ml microcentrifugation tube.

### Sample Preparation for LC-MS/MS Analysis and Data Acquisition

For proteomic analysis, the tissue specimens were prepared as previously described using the direct trypsinization protocol (DTR)^21^. Briefly, protein extraction was performed by adding 100 μl of buffer containing 0.1 % Rapigest in 0.1 M HEPES at pH 8.0 to each tissue sample. Tissue homogenization was performed using sonification in a Bioruptor (Diagenode) (10 cycles with 50/10 sec on/off) followed by the heat-induced antigen retrieval (HIAR) incubating the samples for 2 h at 95°C. Protein concentration of the supernatant was measured using the BCA assay (ThermoScientific) and 100 μg of Protein for each sample was reduced by incubating with 5 mM Dithiothreitol (DTT) for 15 min and alkylated by incubating with 15 mM Iodoacetamide (IAM) for 15 min in the dark. A two-step protein digestion was performed by adding 2 μg of Trypsin and incubating for 2 h at 50°C followed by adding another 2 μg of Trypsin and incubation at 37°C overnight^33^. After digestion, samples were acidified by adding Trifluoroacetic acid (TFA) to a final concentration of 2 % and incubating at 37°C for 30 min. For peptide clean-up, mixed-phase columns (PreOmics) were applied according to the manufacturer’s protocol^34^. Following BCA measurement, 4 μg of peptides of each individual sample were transferred to fresh tubes, vacuum dried, and stored at −80°C until LC-MS/MS measurement.

### LC-MS/MS Data Acquisition and Analysis

For LC-MS/MS analysis 300 ng per sample were measured using an Orbitrap Q-Exactive plus (Thermo Scientific) mass spectrometer coupled to an Easy nanoLC 1000 (Thermo Scientific) with a flow rate of 300 nl/min. Buffer A contained 0.3 % acetic acid in water and buffer B 0.3 % acetic acid in 80 % acetonitrile. Peptides were separated with an increasing gradient of organic solvent (0-60 % acetonitrile in 90 min) on an analytical column (Acclaim PepMap column (Thermo Scientific), 2 μm particle size, 100 Å pore size, length 150 mm, inner diameter 50 μm). The MS was operated in a data-dependent mode and each MS scan was followed by a maximum of ten MS/MS scans.

For data analysis, the MaxQuant (V1.6.0.16) software was used with a reviewed human database (retrieved from UniProt, October 6, 2017) containing 20,188 sequences^35^. Additionally, eleven synthetic peptides (iRT peptides) were added to the database. Decoys for database search were generated using the revert function. Precursor, main search, and fragment mass tolerance were set to be 20, 4.5, and 20 ppm, respectively. The peptide search included a fixed modification of carbamidomethyl cysteine as well as the oxidation of methionine and the acetylation of the protein (n-term) as variable modifications. Tryptic cleavage specificity with up to two missed cleavages was used with a minimum peptide length of seven amino acids. The false discovery rate (FDR) for peptides and proteins was set to 0.01. Files obtained by MaxQuant were processed using the open-source statistical software package R (V4.0.2). Decoy sequences and potential contaminant entries were removed prior to statistical analysis. Raw intensities were log2 transformed and statistical inference of differentially regulated peptides was performed using the limma package (V.3.44.3)^36^ Reported P-values were corrected at a Benjamini-Hochberg FDR of 0.05^37^. Further analyses were performed using mixOmics package (V6.12.2) (hierarchical clustering, clustering distance: Euclidean distance, PCA), corrPlot package (V0.84) and EnhancedVolcano package (V1.6.0)^38,39^. Gene ontology (GO) and REACTOME enrichment analysis was performed using the topGO package (V2.40.0) and the ReactomePA package (V1.32.0)^40^.

### Immunohistochemistry (IHC)

Immunohistochemical staining of ALDH1A1, ALDOB, and DPP4 was performed as previously described using specific antibodies mouse anti-human ALDH1A1 (R&D, MAB5869), rabbit anti-human ALDOB (AbCam, ab75751), and mouse anti-human DPP4 (AbCam, ab114033)^26^. Briefly, 2 μm tissue slices were deparaffinized and heat-induced antigen retrieval (HIAR) was performed. Tissue sections were stained by applying the following steps: incubating with 10 % H2O2 for 5 min, with primary antibody for 30 min, with secondary antibody for 10 min, and with horseradish peroxidase for 20 min, and lastly with 3,3’-diaminobenzidine for 10 min. Between each of the aforementioned steps, the tissue was rinsed using a washing buffer containing 50 mM Tris-HCl, 150 mM NaCl, and 0.05% Tween 20. Each tissue slice was counterstained by incubating with hematoxylin for 30 sec and xylene was used as a permanent mounting medium.

### Cell proliferation assay

CaCo-2 and SW480 cells were cultured in DMEM (Gibco) supplemented with 10% foetal calf serum (PAN), 1% penicillin/streptomycin (Gibco), 1% MEM vitamin solution (PAN), 1% MEM non essential amino acid solution (PAN), and 1% L-glutamine (Gibco). Prior to inhibitor treatment 1 ×10^5^ cells/well were incubated in 6-well plates for 24 h. For the inhibition of Argininosuccinate synthase (ASS1) cells were incubated for 48 h after adding 5 mM (final conc.) of N-methyl-DL-aspartic acid (MDLA) (Santa Cruz), solved in water, or an equal volume of water as a control. For the inhibition of Thymidylate kinase (DTYMK) cells were incubated for 72 h after adding 2 μM (final conc.) of the YMU-1 compound (Sigma-Aldrich), solved in 15 % DMSO, or an equal volume of 15 % DMSO as a control. Following the YMU-1 treatment, the cells were incubated for 4 h after adding either the FOLFOX regimen (50 μg/ml 5-fluorouracil, 40μg/ml oxaliplatin, and 10μg/ml folinic acid) or an equal volume of 15 % DMSO. Subsequently, the cell culture medium was exchanged and cells were incubated in fresh DMEM including supplements for 24 h. Adherent cells for both inhibitor treatments were washed using PBS (Gibco) and harvested by trypsinization. Cell proliferation was determined by counting with Trypan Blue staining using EVE™Automated Cell Counter (NanoEntek) according to the manufacturer’s protocol.

## Results and Discussion

### A straightforward direct trypsinization protocol facilitates the proteomic investigation of patient-derived FFPE tissue specimens

We aimed for a comprehensive proteome investigation of primary colorectal carcinoma (CRC) and their derived liver metastasis in a small cohort of seven CRC patients (**Figure 1A**). The tissue was formalin-fixed and paraffin-embedded (FFPE) following surgical removal. The FFPE procedure preserves cellular as well as tissue morphology and prevents tissue degradation. Thus, FFPE specimens are a standard for histopathological diagnostics and can be stored for decades in vast tissue archives. FFPE specimens provide a valuable resource for proteomic investigations of a variety of malignancies^25–28^. Here we used 10 μm thin tissue slices, which were first deparaffinized and subsequently used for tissue macro dissection by an experienced pathologist, focusing on tumorous tissue of either the primary CRC or the resulting liver metastasis (**Figure 1B**). A direct trypsinization protocol was applied using the macro dissected tumorous tissue^21^. Of note, the samples were prepared by students under the supervision of experienced researchers as part of a two-day practical course, emphasizing the straightforwardness of the applied protocol. The entire proteomic workflow is illustrated in **Figure 1**.

**Figure 1.**
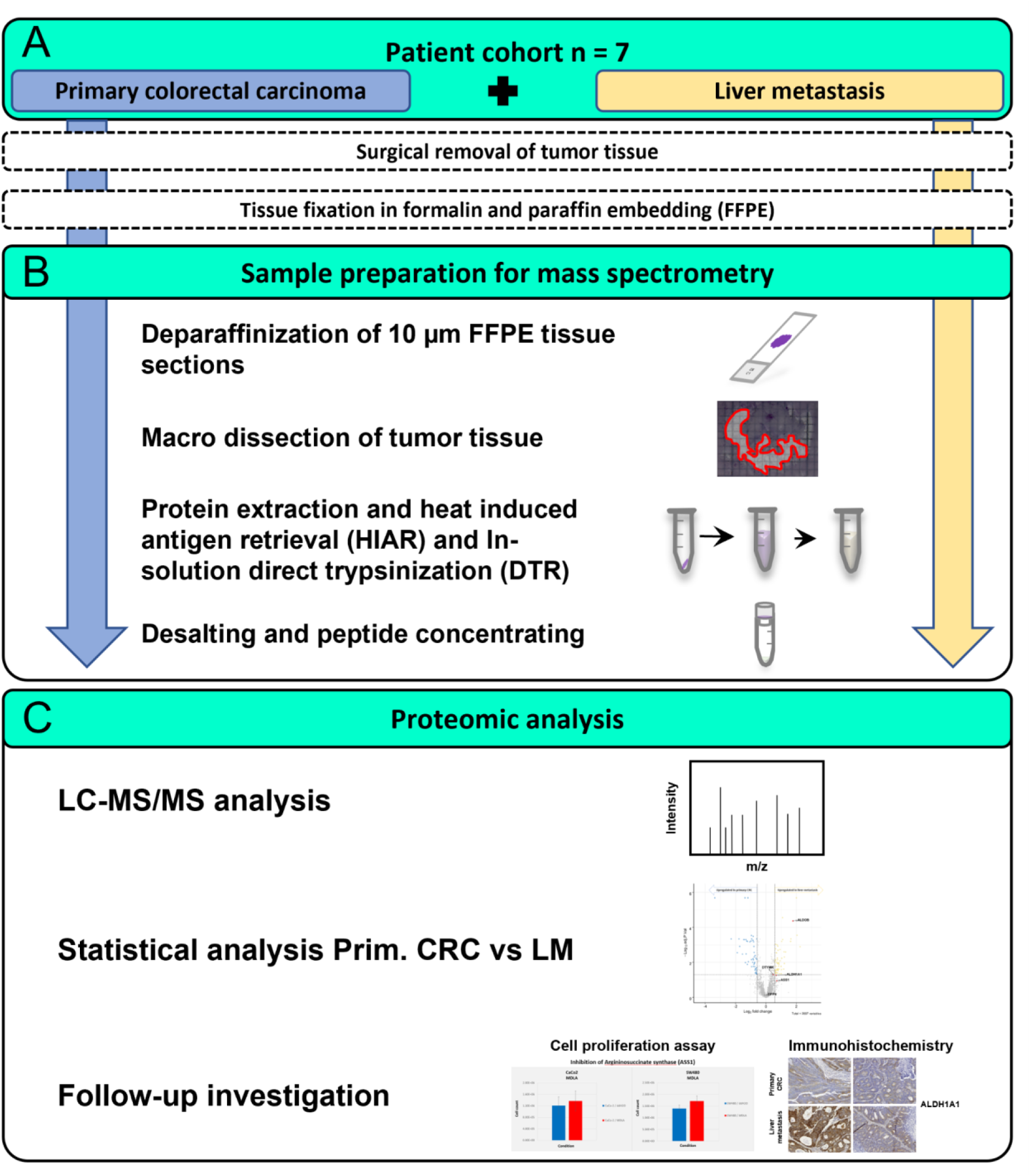
Overview of the experimental design to investigate tissue of primary colorectal carcinoma and patient-matched liver metastasis. (A) Primary colorectal carcinoma tissue (blue) and patient-matched liver metastasis tissue (yellow) were collected from seven patients. Following surgical removal, the tissue was conserved using formalin fixation and paraffin-embedding (FFPE). (B) Small, 10 μm thick tissue slices were deparaffinized. For each sample, the tumorous tissue was macro dissected and transferred to fresh microcentrifugation tubes. Protein extraction and heat-induced antigen retrieval (HIAR) in combination with a direct trypsinization (DTR) protocol was applied. Following peptide clean-up, the samples were measured using liquid chromatography-tandem mass spectrometry (LC-MS/MS). Differential expression analysis was performed and individual target proteins were used for Immunohistochemistry (IHC) and cell proliferation assays.

All patients developed distant metastasis in the liver and/or the peritoneum (**Table 1**). Further details including the tumor localization, the grading, and the TNM classification are summarized in **Table 1**. Generally, patients were between 48 and 75 years old, including two female and five male patients. Patient data and raw LC-MS/MS data have been deposited in the European Genome-phenome Archive (EGA) and can be accessed via a Data Access Committee (DAC).

### A small cohort of primary CRC and liver metastasis FFPE tissue allows robust and reproducible proteome investigation

All samples (n = 7 primary CRC and n = 7 liver metastasis) were macro dissected, prepared, and measured in duplicates. Protein intensities of the two replicates for each patient sample showed a high correlation with Pearson correlation coefficients ranging from 0.74 – 0.93 (**Supplementary Figure 1**). Proteins were included for further analysis if they have been quantified in at least one replicate per duplicate; wherever possible, a mean intensity per duplicate was calculated. Hereby, more than 2,400 proteins were identified and quantified in all of the liver metastasis samples. Considering the CRC and in five out of seven primary CRC samples (**Figure 2**). Interestingly, we identified higher numbers of proteins in the CRC-LM (on average 2,606) as compared to the primary CRC samples (on average 2,383).

**Figure 2.**
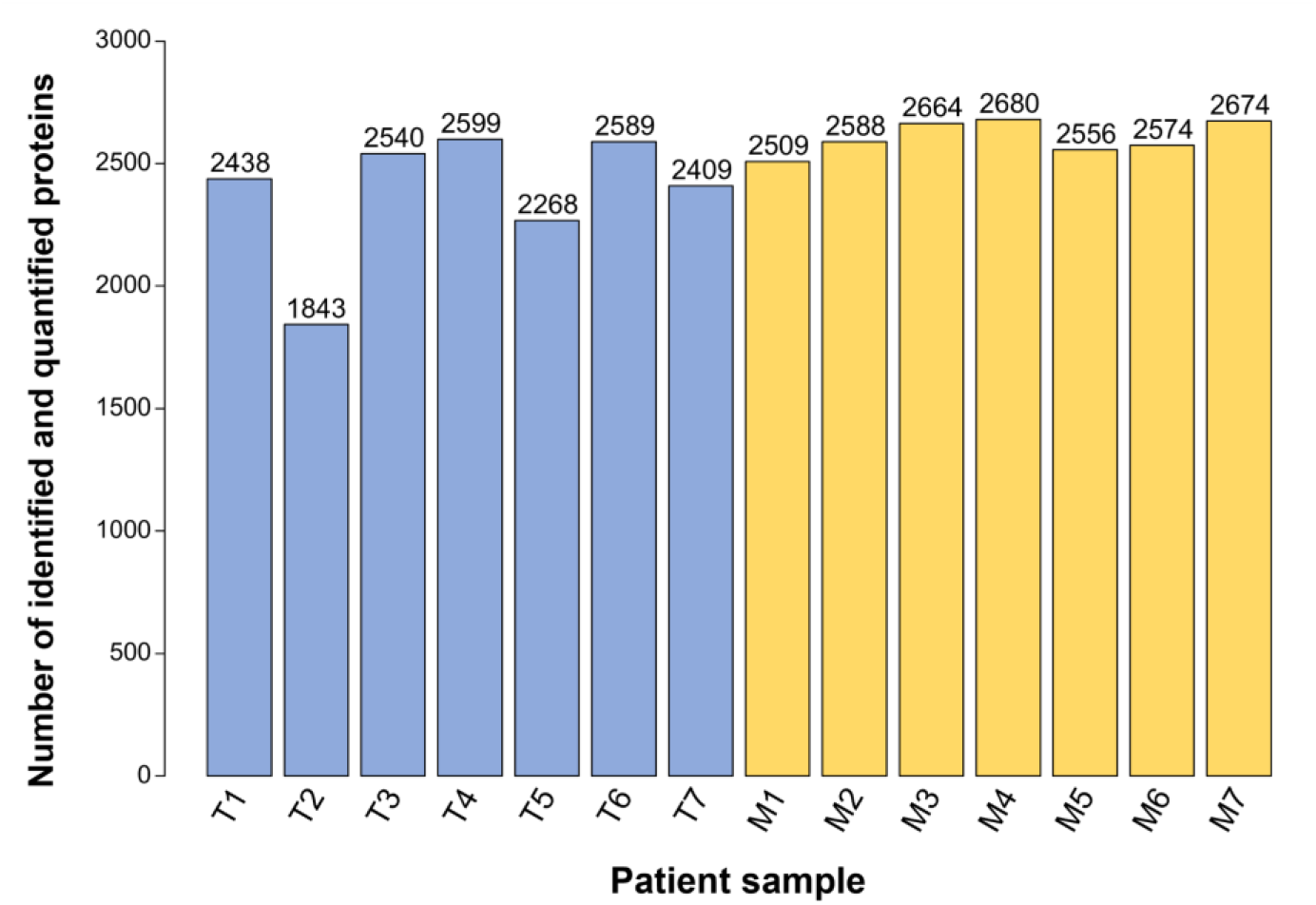
Overview of identified and quantified proteins in primary colorectal carcinoma and patient-matched liver metastasis. The bar chart shows the number of identified and quantified proteins in primary colorectal cancer (T, blue) and liver metastases (M, yellow) samples from n = 7 patients. As described in the material and methods section, two technical replicates per sample were conducted; shown here are identified and quantified in at least one of the measurements.

Unsupervised principal component analysis (PCA) shows a clear separation of the primary CRC samples and the corresponding liver metastases (**Figure 3A**). The liver metastases samples cluster together whereas there are two outliers in the primary CRC samples. The similarity of the samples from either the primary CRC or the liver metastases samples is further highlighted by the formation of two distinct clusters in the hierarchical clustering analysis (**Figure 3B**). Remarkably, the samples cluster according to the tumorous tissue origin, rather than according to the individual patients. This is of particular importance since the primary-metastasis pairs were patient-matched; hence emphasizing truly distinct proteome biology that extends beyond inter-patient heterogeneity even within the comparably small cohort size.

**Figure 3.**
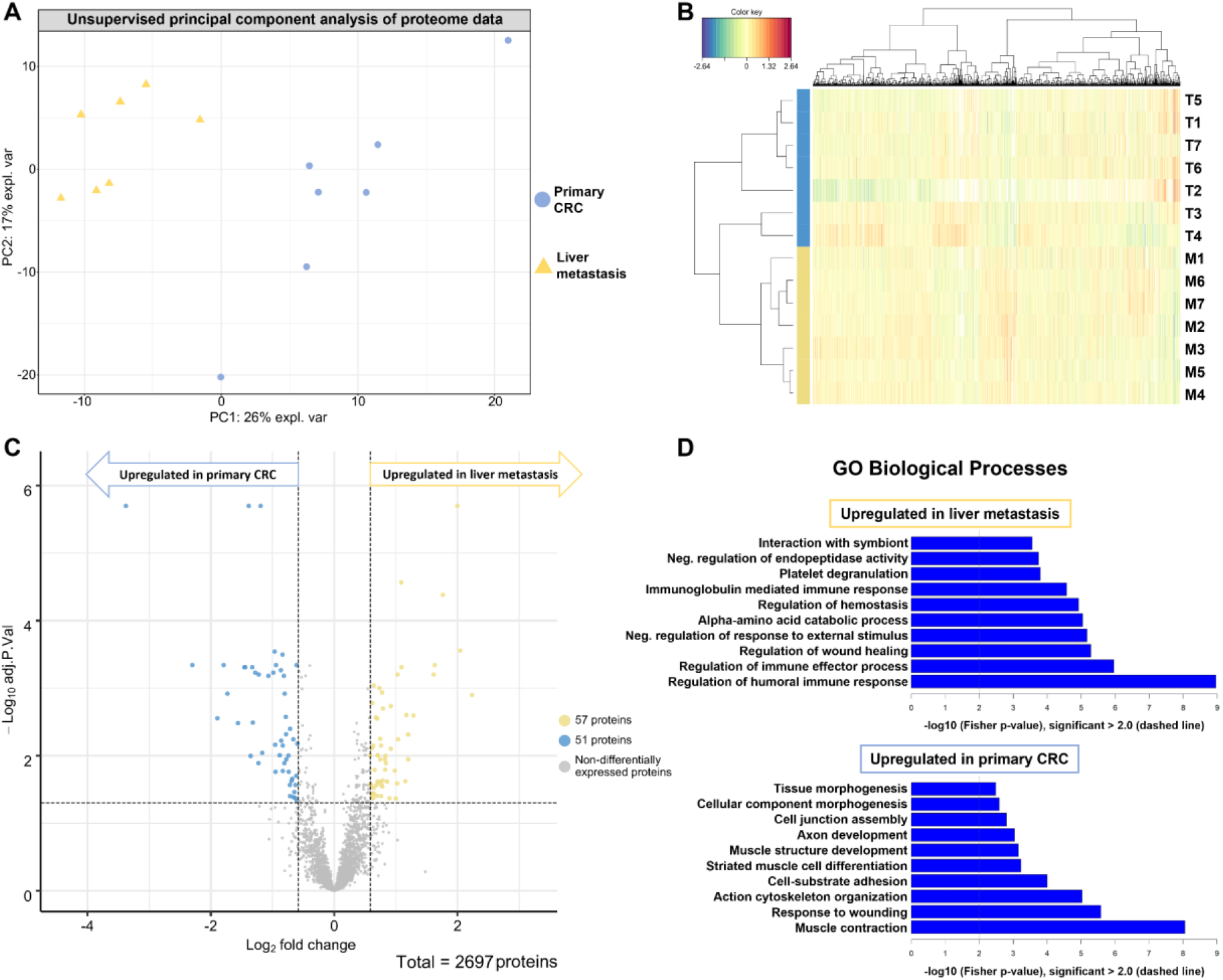
Unsupervised and statistical analysis of primary CRC and liver metastasis. The average protein intensity of the two replicates per patient for the primary colorectal carcinoma (T, blue) and the liver metastasis (M, yellow) was computed. Proteins that were at least qualified in 4 out of the 14 samples were used for unsupervised principal component analysis (A) and hierarchical clustering (B). (C) Volcano plot showing proteins with their respective −log10 adjusted p-value and the log2 fold change for the comparison of liver metastases against primary CRC tissue from seven patients. Of the 2697 proteins, 57 were significantly more abundant in liver metastasis, whereas 51 proteins were significantly more abundant in primary colorectal tumors (adjusted p-value < 0.05). (D) Gene ontology (GO) analysis of the significantly dysregulated proteins shows upregulated biological processes for each tumor tissue.

### Statistical analysis reveals differentially expressed proteins and distinct proteome biology in primary CRC and corresponding liver metastases

To identify significantly dysregulated proteins between the primary CRC and the CRC-LM a statistical analysis using linear models of microarray analysis (limma) was performed^36^. As criteria for significant changes, we requested an adjusted p-value < 0.05 as well as a change in abundance (fold change) above 1.5 for proteins that were upregulated in liver metastasis and below 1.5 for proteins that were upregulated in the primary CRC (**Figure 3C**); corresponding to an increase or decrease in abundance of at least 50 %. In total, we identified 108 significantly dysregulated proteins, among which 57 proteins were enriched and 51 proteins were depleted in the CRC-LM as compared to the CRC (**Table 2**). As expected, many of the upregulated proteins in liver metastasis are involved in metabolic processes such as gluconeogenesis and fructose metabolism, e.g. the pyruvate carboxylase and the fructose-bisphosphate aldolase B as well as Fructose-1,6-bisphosphatase 1 (**Table 2**). Interestingly, we notice enrichment of multiple members of the complement system in the liver metastasis, including proteins linked to complement components C1, C4, C5, and C9. On the other hand, multiple proteins associated with muscle contraction and cell junction assembly are depleted in the liver metastases, including Desmin and Synemin as well as Filamin-C (**Table 2**).

**Table 2.**
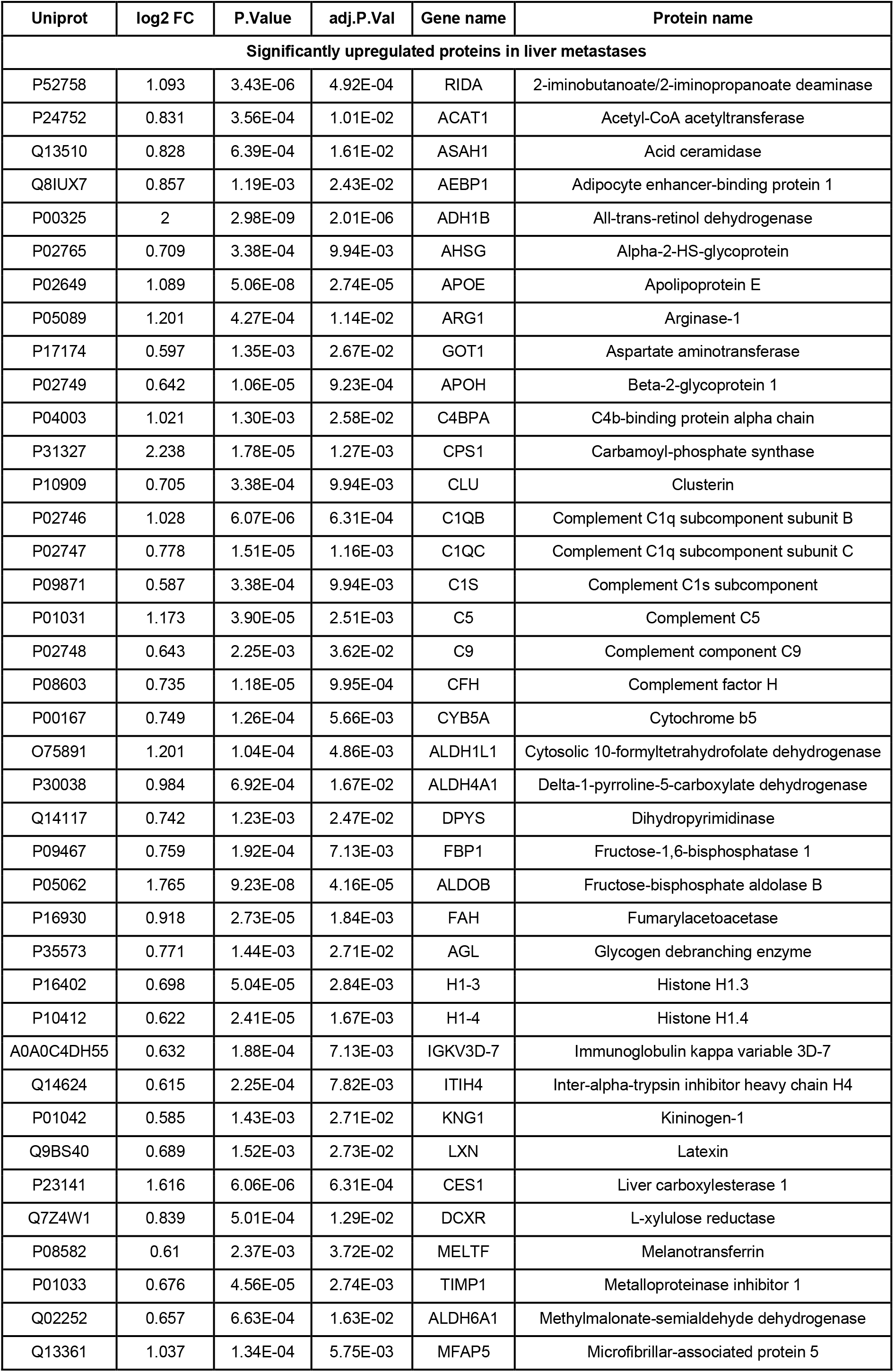
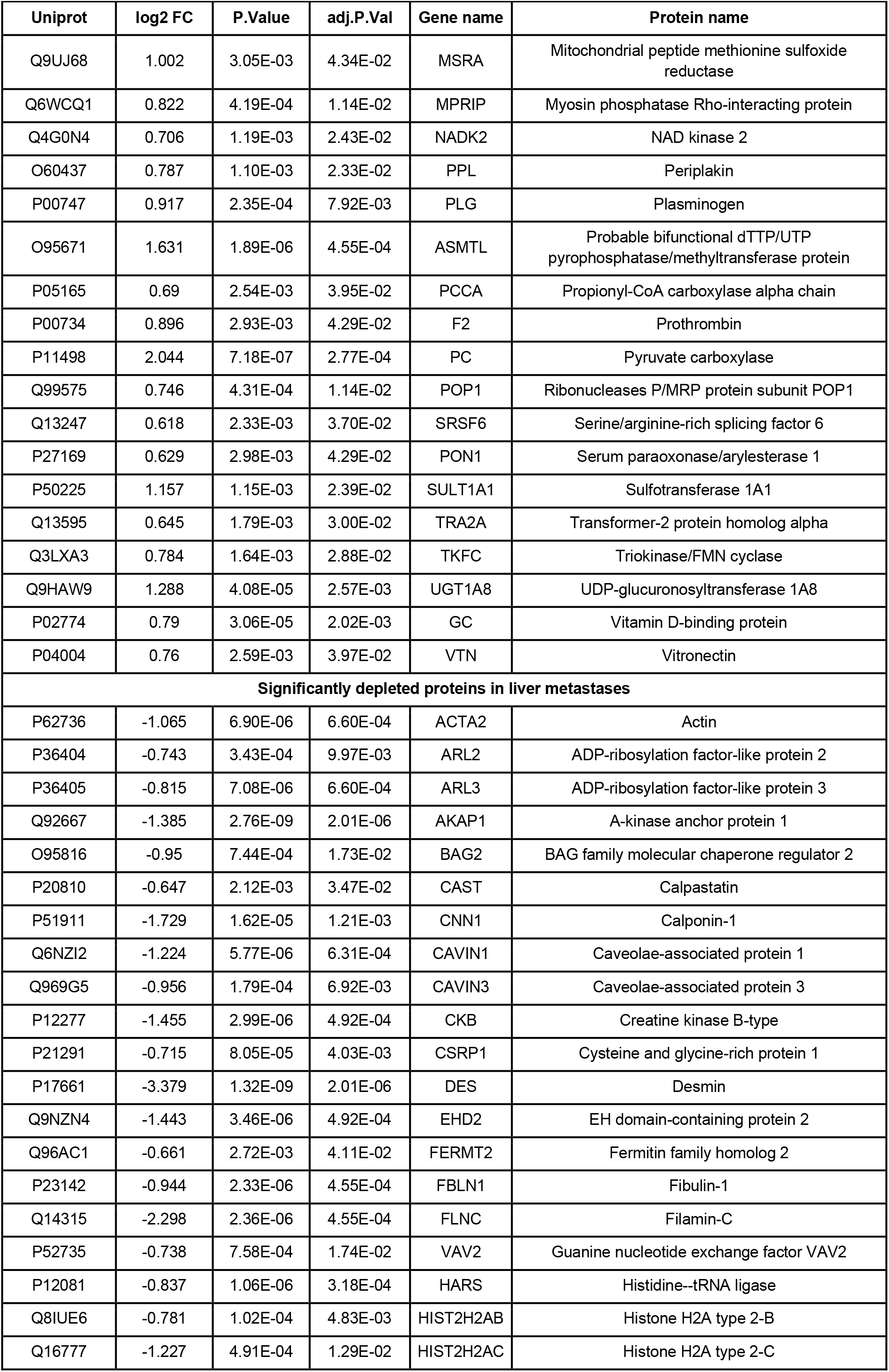
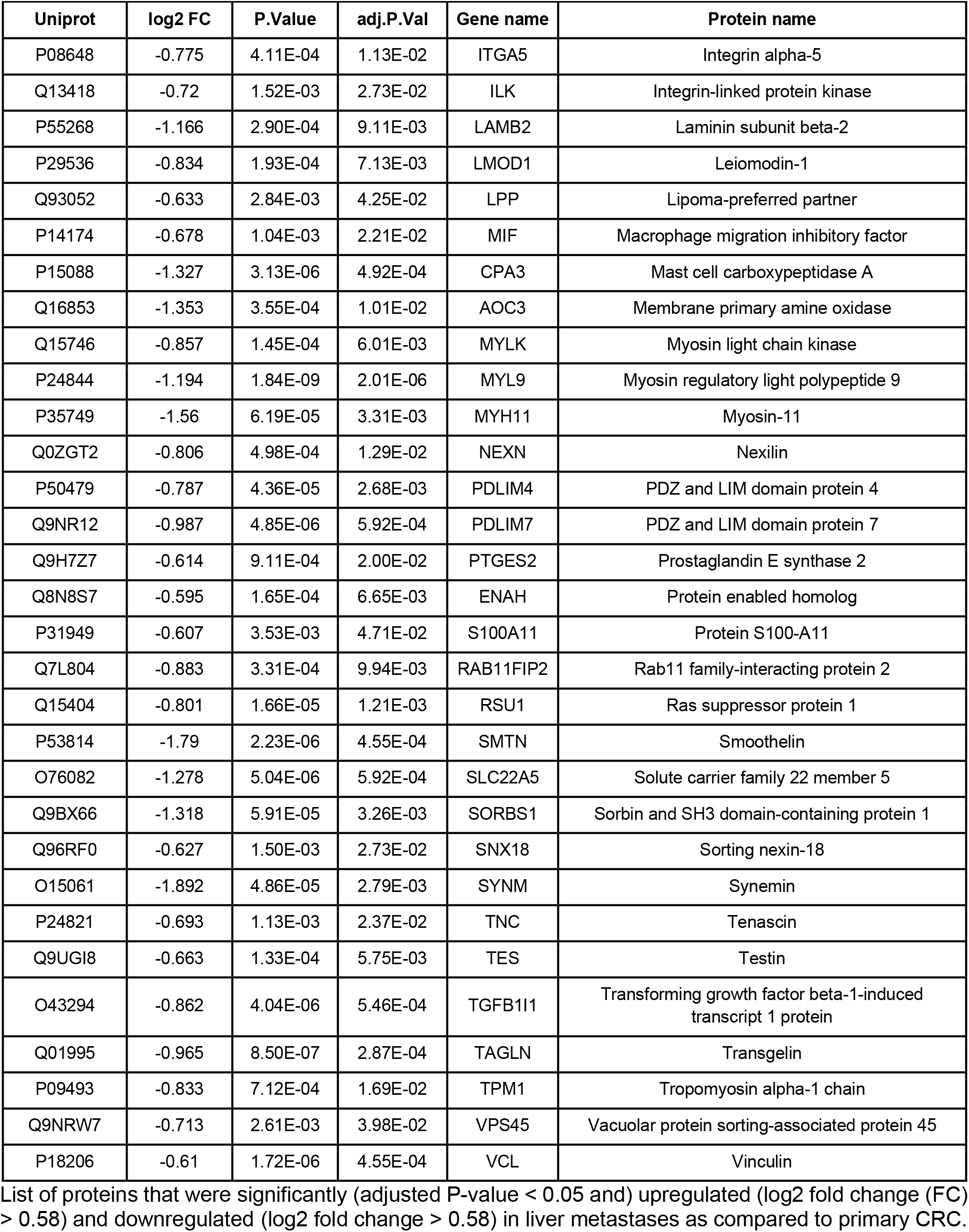
List of significantly dysregulated proteins in liver metastases as compared to primary CRC.

### CRC-derived liver metastasis presents an upregulation of biological processes linked to the immune response, whereas primary CRC shows upregulation of structural components

Further, we performed a Gene Ontology (GO) enrichment analysis to identify commonly affected differentially upregulated biological processes in a more systematic manner. To this end, we probed the set of proteins that we found to be either enriched or depleted in the liver metastases as compared to the entirety of identified and quantified proteins (**Figure 3D**).

The protein signature that we found to be enriched in liver metastases mapped to a variety of biological processes associated with metabolic activity. The fingerprint of enriched metabolic processes is interesting. Liver tissue can be expected to be a major source of metabolic enzymes. However, non-malignant liver tissue has been removed by macro dissection prior to proteomic analysis. As will be shown in the section on immunohistochemistry (IHC), metabolic enzymes such as the retinal dehydrogenase (ALDH1A1) and the fructose-bisphosphate aldolase B (ALDOB) are prominently expressed by tumor cells in liver metastases. Several upregulated biological processes in the liver metastasis are associated with the immune response, e.g. the regulation of the immune effector process as well as the regulation of the humoral immune response. Moreover, proteins associated with the negative regulation of endopeptidase activity and the regulation of wound healing are higher expressed in liver metastasis than in primary CRC tissue. The protein signature that we found to be depleted in liver metastases mapped mainly to structural biological processes such as the actin cytoskeleton organization, the cell junction assembly as well as muscle contraction (**Figure 3D**). Those results indicated a more active immune response within the metastatic tissue as compared to the primary tumor location.

### Immunohistochemistry reveals tumor-cell expression of metabolic proteins primary CRC and their corresponding liver metastasis

To investigate the cell-type-specific expression and spatial distribution of selected proteins of interest we used immunohistochemistry (IHC) (see **Figure 4** and **Supplementary Figure 6**). Sufficient sample material for IHC was available for five pairs of primary and metastatic tissue and it was not possible to include further specimens. The staining of ALDH1A1 highlighted its expression in tumorous tissue in both the primary CRC as well as the liver metastasis (**Figure 4A**). Furthermore, we observe in three cases a stronger expression of ALDH1A1 in the tumor cells of the liver metastases as compared to the primary CRC, consistent with the observed upregulation in the proteome data (**Supplementary Figure 6**). IHC staining of ALDOB failed to produce a strong signal in most cases (**Figure 4B**). Nevertheless, expression of ALDOB in tumorous tissue of primary CRC as well as liver metastasis was detected in most of the assessed samples. Interestingly, we observe a prominent upregulation of ALDOB in the CRC-derived liver metastases in the proteome data, which has also been described in previous studies (**Supplementary Figure 6**)^41^. Furthermore, overexpression of ALDOB has been associated with poor prognosis, promoting tumor progression^42^. We have also probed for Dipeptidyl peptidase 4 (DPP4) as a protein that showed a rather consistent abundance (log2 FC = −0.12; adjusted p-value = 0.73) between the primary CRC samples and the liver metastases (**Supplementary Figure 6**). For DPP4 very prominent staining in most of the cases is evident, without any visible alteration between the two tumor entities (**Figure 4C**).

**Figure 4.**
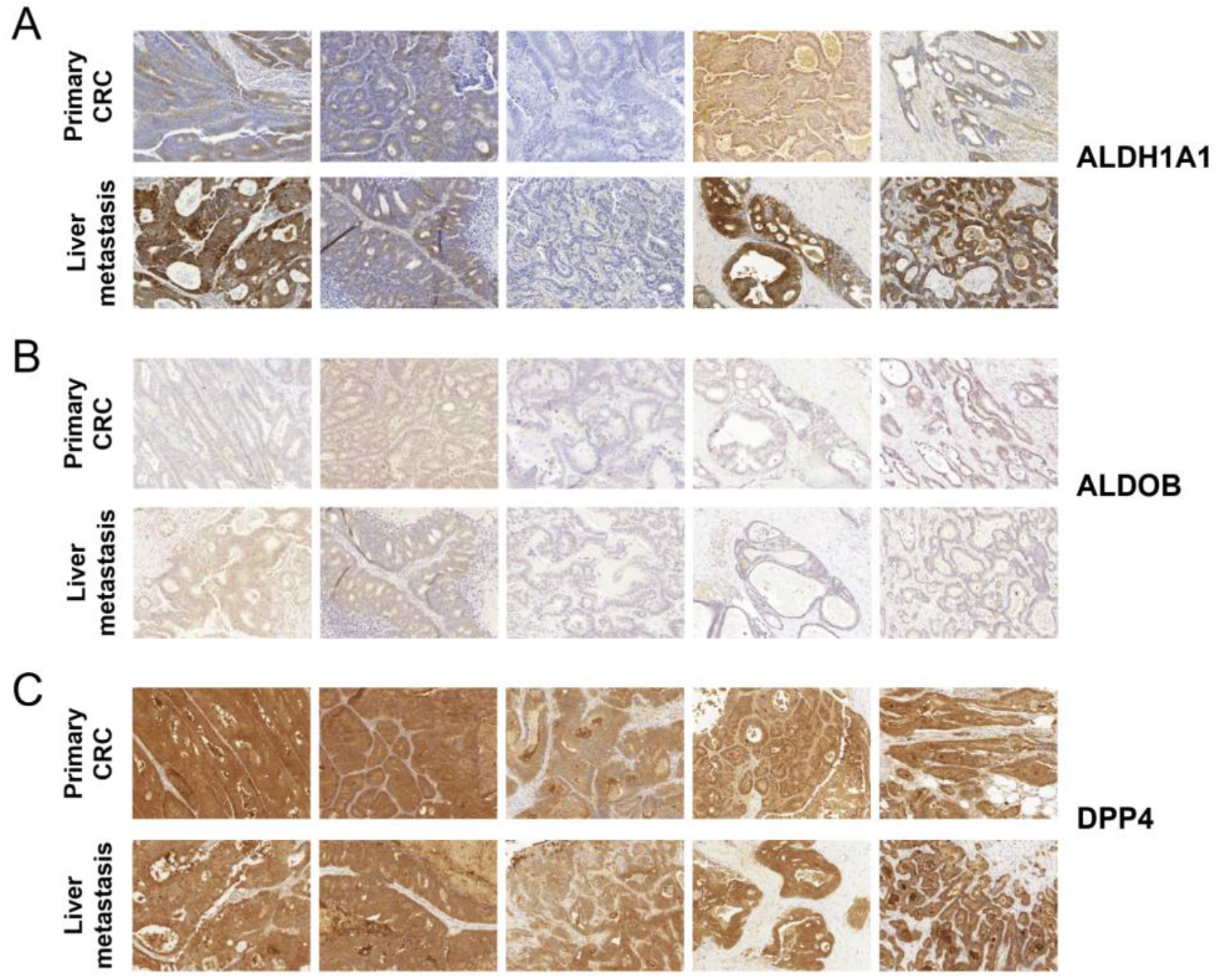
Follow-up investigation using Immunohistochemistry (IHC) on primary colorectal carcinoma (CRC) and liver metastasis tissue of 5 representative patients. IHC staining for A) ALDH1A1, B) ALDOB and C) DPP4.

### Functional follow-up investigation yields a more comprehensive understanding of CRC biology by inhibition of two metabolic proteins that were enriched in liver metastases

A cell proliferation assay was performed to investigate the inhibition of two metabolic proteins that were upregulated in liver metastases using two established colorectal cancer cell lines SW480 and CaCo-2. Inhibition of Argininosuccinate synthase (ASS1) has been described to reduce levels of the oncogenic metabolite fumarate and results in impaired proliferation in SW620 cells^43^. The inhibition of ASS1 using N-methyl-DL-aspartic acid (MDLA) does not lead to a decrease in proliferation in either SW480 or CaCo-2 cells as compared to the control (**Figure 5A**). Furthermore, the inhibition of Thymidylate Kinase (DTYMK) has been shown to sensitize tumor cells to doxorubicin treatment in vitro and in vivo^44^. Here, two colorectal cell lines SW480 and CaCo-2 were treated with the established chemotherapeutic regimen FOLFOX^45^. In both cell lines, a clear decrease in cell proliferation upon chemotherapeutic treatment can be observed (**Figure 5B**). However, the previously described chemosensitization effect following the inhibition of DTYMK in combination with doxorubicin treatment is not detectable in neither SW480 nor CaCo-2 cells using the FOLFOX regimen.

**Figure 5.**
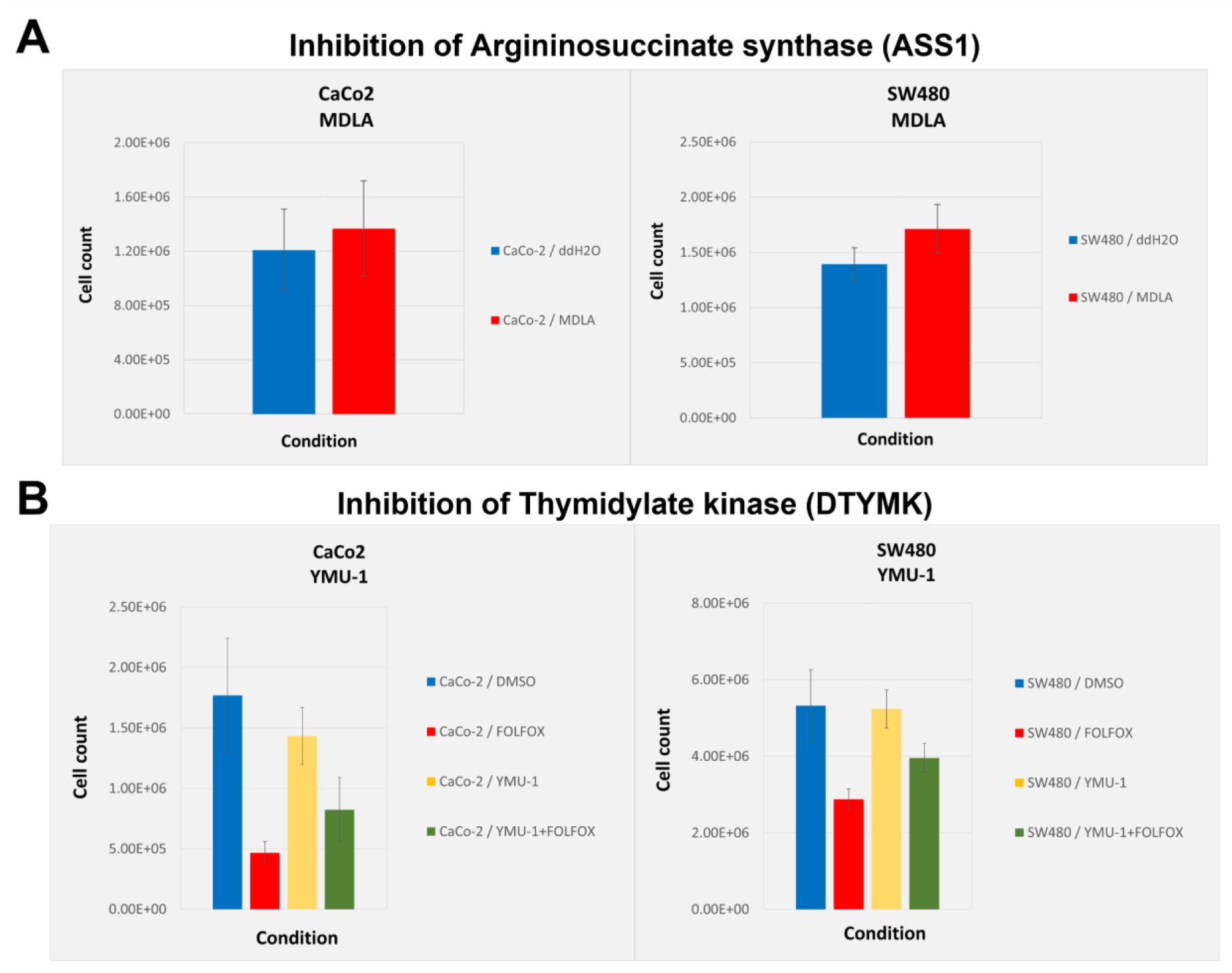
Functional follow-up analysis of proteins that were upregulated in liver metastasis using two colorectal cell lines. Proliferation assays were performed using two established colorectal cancer cell lines CaCo-2 and SW480. Bar plots show the mean cell count of five biological replicates per condition, including the standard deviation as error bars. A) Bar plot showing the cell count following the inhibition of Argininosuccinate synthase (ASS1) using N-methyl-DL-aspartic acid (MDLA). Cells treated with water (blue) or with 5mM MDLA (red). B) Bar plot showing the cell count following the inhibition of Thymidylate kinase (DTYMK) using the YMU-1 compound. Cells were either treated with 15% DMSO (blue), with the FOLFOX regimen (fluorouracil, oxaliplatin, and folinic acid) (red), with 2μm YMU-1 (yellow) or a combination of YMU-1 and FOLFOX (green).

### Tissue macro dissection improves the sensitivity of proteome investigation of primary CRC and their resulting liver metastasis

To investigate the necessity and the benefits of the tissue macro dissection we have also prepared adjacent FFPE slices without prior dissection of the tumorous tissue. The numbers of identified and quantified proteins in each sample are comparable to the macro dissected approach (**Supplementary Figure 2**). It appears that in the non-dissected tissue, higher numbers of proteins are identified in primary CRC samples as compared to the liver metastasis samples. Unsupervised PCA and hierarchical clustering show a clear separation of the primary CRC and the liver metastasis in the non-dissected tissue samples (**Supplementary Figure 3**). Noteworthy, the primary CRC from patient 4 is close to the liver metastasis from patient 1 in both the non-dissected as well as the dissected tissue analysis (see **Figure 3B** and **Supplementary Figure 3B**). In the non-dissected comparison of primary CRC and liver metastasis over 200 proteins are significantly dysregulated (**Supplementary Figure 4A**). However, the GO enrichment analysis reveals mostly biological processes associated with metabolism as upregulated in the liver metastases samples (**Supplementary Figure 4B**). On the other hand, GO enrichment analysis of proteins depleted in the (non-dissected) liver metastases samples show similar biological processes including cell junction assembly and cytoskeleton organization. Only in the macro dissection approach, the upregulation of proteins associated with the immune response becomes apparent (see **Figure 3D** and **Supplementary Figure 4**). Thus, (and as expected) the tissue macro dissection improves the sensitivity of proteome investigations using primary CRC and liver metastasis tissue. The improved sensitivity and thus the benefits of a tissue macro dissection are also highlighted in a REACTOME analysis of the differentially expressed proteins in the dissected and the non-dissected proteome investigation (**Supplementary Figure 5**). Here, the active immune response and the activation of the complement system are only observable in the dissected approach, whereas only metabolism-associated processes are identified in the non-dissected approach.

## Conclusion

We show the straightforwardness and practicability of a reproducible and robust protocol for the detailed proteome investigation using patient-derived FFPE tissue. In this small cohort study, the distant metastasis clearly separates away from the primary tumor strongly suggesting a prominent difference in proteome biology. Furthermore, the relatively small number of patient samples (n = 7) in combination with the tissue dissection enabled the identification of previously described Fructose-bisphosphate aldolase B (ALDOB) as specifically enriched in CRC-derived liver metastatic tumor tissue^41,42^. Despite a prominent metabolic molecular fingerprint, we were able to detect an enhanced immune response in the liver metastases as compared to the primary CRC. Detailed follow-up investigation showed tumor-cell expression of Retinal dehydrogenase 1 (ALDH1A1) that has been described in various tumor-related contexts, with prognostic characteristics in breast, pancreatic, and prostate cancer^25,46–49^. Here, a functional follow-up investigation in two established CRC cell lines does not show a previously described chemosensitization effect upon inhibition of the Thymidylate kinase^44^. Even though such a chemosensitization has been described in other cancer cell lines with doxorubicin treatment, we feel that using the FOLFOX regimen might provide additional insights, since this is one of the currently applied treatments in patients. This project illustrates the potential and added value towards a more comprehensive understanding of clinical malignancies using detailed proteome investigations, even in smaller cohorts. Proteome-wide studies in larger CRC cohorts are needed to elucidate further on the molecular mechanisms of metastasis formation as well as tumor progression.

## Additional Files

**Supplementary Figure 1. Correlation analysis of technical replicates of primary colorectal carcinoma and liver metastasis samples from seven patients.** Correlation analysis of the measured intensities of primary colorectal carcinoma (A) and liver metastasis (B) tissue. Pearson correlation coefficients are shown in the bottom left corner and illustrated as ellipses in the upper right corner. Correlation for the two technical replicates of the same patient sample is highlighted in green.

**Supplementary Figure 2. Overview of identified and quantified proteins in non-dissected primary CRC and liver metastases.**

The bar chart shows the number of identified and quantified proteins in primary colorectal cancer (blue) and liver metastases (yellow) samples from n = 7 patients.

**Supplementary Figure 3. Unsupervised analysis of non-dissected primary CRC and liver metastases.**

Proteins that were at least qualified in 4 out of the 14 samples were used for unsupervised principal component analysis (A) and hierarchical clustering (B).

**Supplementary Figure 4. Statistical analysis of non-dissected primary CRC and liver metastases of n = 7 patients.**

A) Volcano plot showing proteins with their respective −log10 adjusted p-value and the log2 fold change for the comparison of liver metastases against primary CRC tissue from seven patients. Of the 2461 proteins, 105 were significantly more abundant in liver metastases, whereas 100 proteins were significantly more abundant in primary CRC (adjusted p-value < 0.05). B) Gene ontology (GO) analysis of the significantly dysregulated proteins shows upregulated biological processes for each tumor tissue.

**Supplementary Figure 5. REACTOME analysis of primary CRC and liver metastases of n = 7 patients.**

REACTOME analysis of the significantly dysregulated proteins in A) the dissected and B) the non-dissected proteome data. The dot plot shows the affected pathways and the number of dysregulated genes as well as the adjusted p-value.

**Supplementary Figure 6. Statistical analysis highlighting proteins of interest in the comparison of primary CRC and their corresponding liver metastases.** Volcano plot showing proteins with their respective −log10 adjusted p-value and the log2 fold change for the comparison of liver metastases against primary CRC tissue from seven patients. Of the 2697 proteins, 57 were significantly more abundant in liver metastasis, whereas 51 proteins were significantly more abundant in primary colorectal tumors (adjusted p-value < 0.05). Proteins of interest (POI) that were used for follow-up investigation using either Immunohistochemistry (IHC) or proliferation assays are highlighted in red.

## Acknowledgment

OS acknowledges funding by the Deutsche Forschungsgemeinschaft (DFG, SCHI 871/17-1, NY 90/6-1; SCHI 871/15-1, GR 4553/5-1, PA 2807/3-1, project-ID 431984000 – SFB 1453 (project P12); project-ID 441891347 – SFB 1479 (project S1); project-ID 423813989/GRK2606 (RTG “ProtPath”), INST 39/766-3 (Z1)), the ERA PerMed programs (BMBF, 01KU1916, 01KU1915A), the German-Israel Foundation (grant no. 1444), and the German Consortium for Translational Cancer Research (project Impro-Rec). SF acknowledges funding by the Deutsche Forschungsgemeinschaft project-ID 89986987 – SFB 850; project-ID 280163318 – FOR 2438. The authors acknowledge the help during proteomic sample preparation from the students that were part of the molecular and cellular biology course (2018/2019). The authors acknowledge the help of Lucas Kook during the analysis of the proteome data.

## Author Contributions

MF conceived the project, performed proteomics, analyzed data, and drafted the manuscript. AJ conceived the project, performed proteomics, provided clinical tissue specimens, and drafted the manuscript. PB performed tissue macro dissection, immunohistochemistry staining, and scoring. OS conceived the project, supervised the proteomics part, and drafted the manuscript. SF supervised the project. All authors have approved the final article.

