## Supplementary figures and images for "Proteome biology of primary colorectal carcinoma and corresponding liver metastases"

### Supplementary Figure 1

#### Slide 1
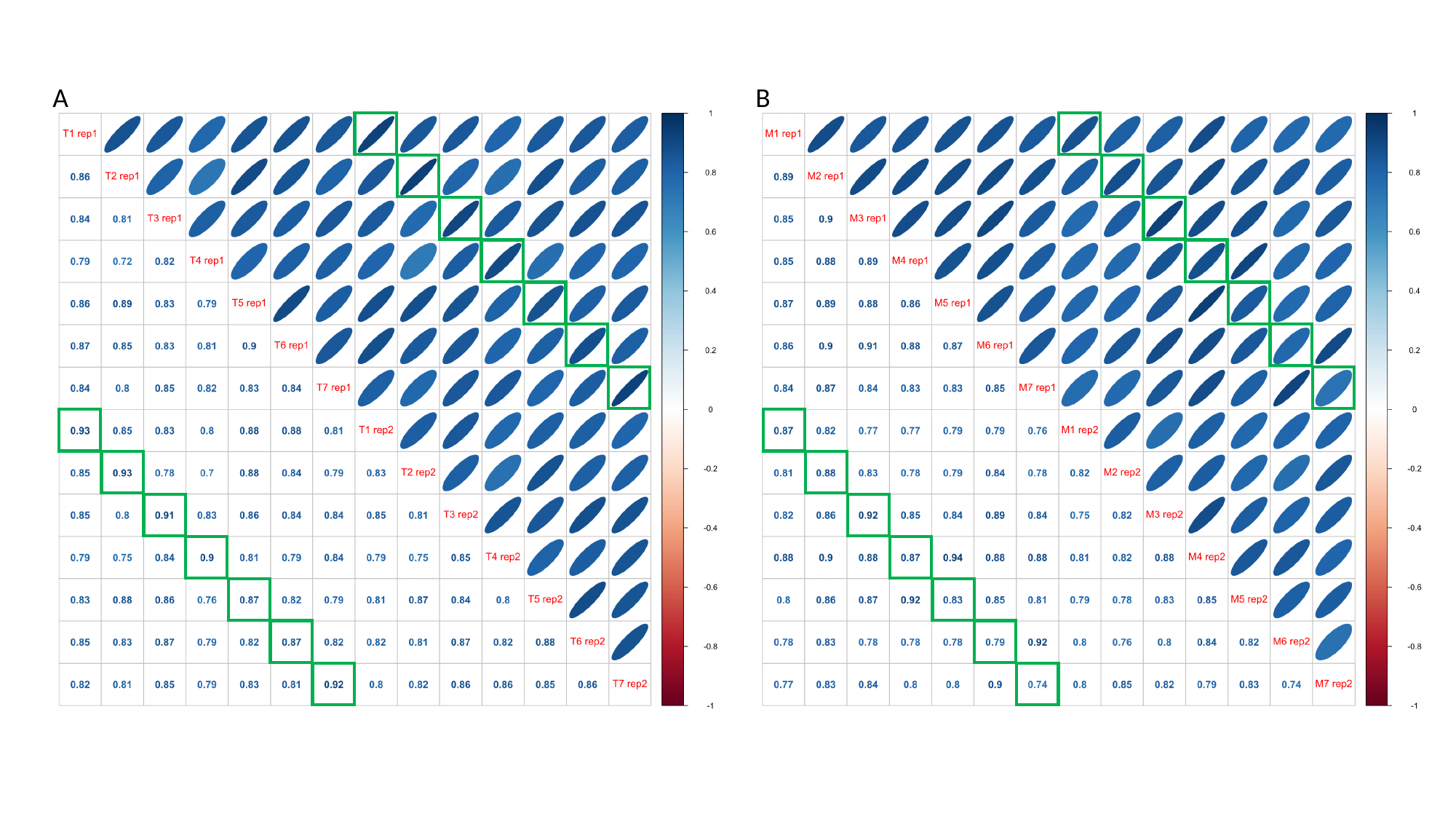

A
B

### Supplementary Figure 2

#### Slide 1
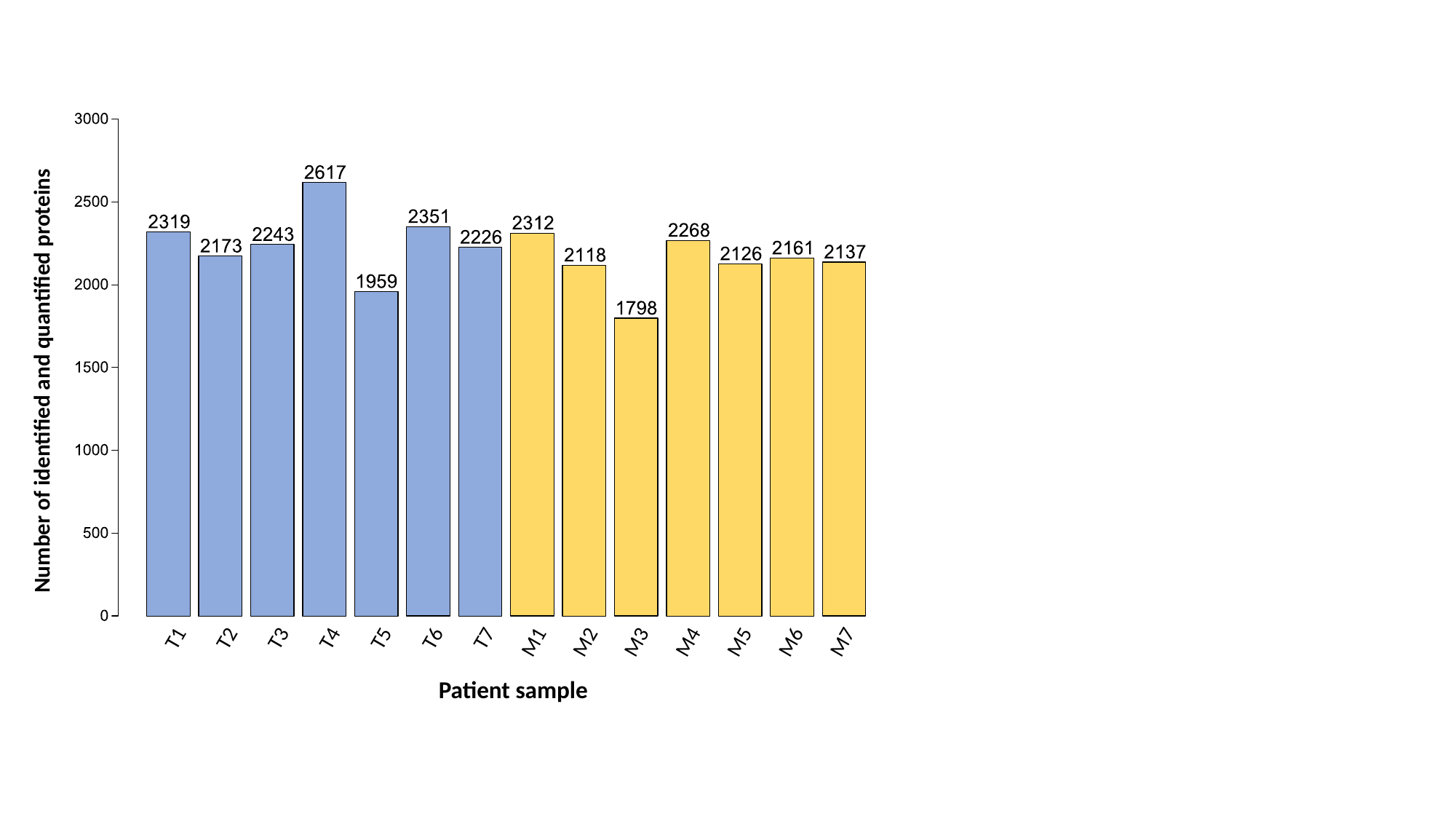

Number of identified and quantified proteins
M7
T1
T2
T3
T4
T5
T6
T7
M1
M2
M3
M4
M5
M6
Patient sample

### Supplementary Figure 6

#### Slide 1
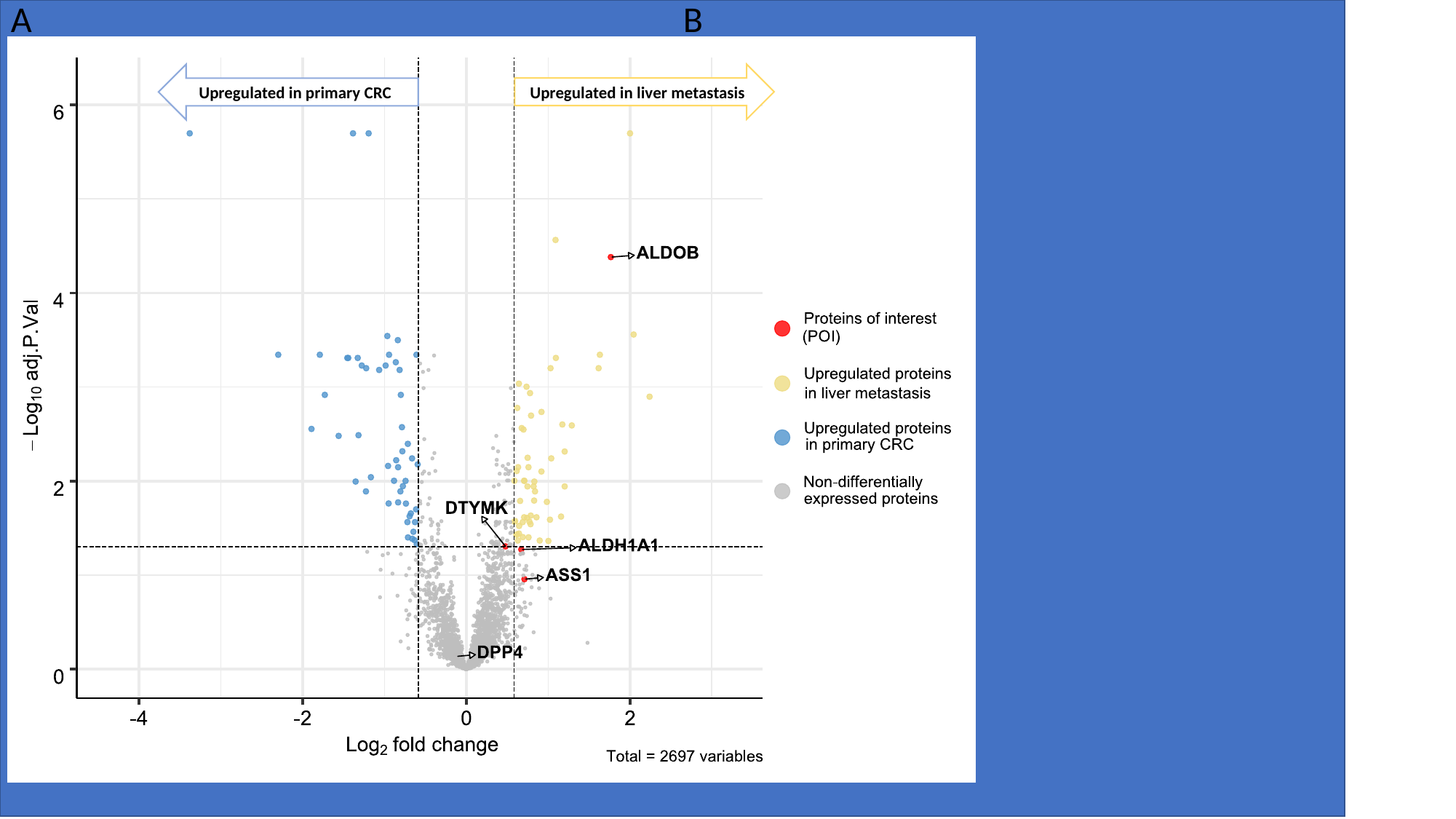

A
B
Upregulated in primary CRC
Upregulated in liver metastasis
