## Supplementary Figure 3 for "Proteome biology of primary colorectal carcinoma and corresponding liver metastases"

### Slide 1
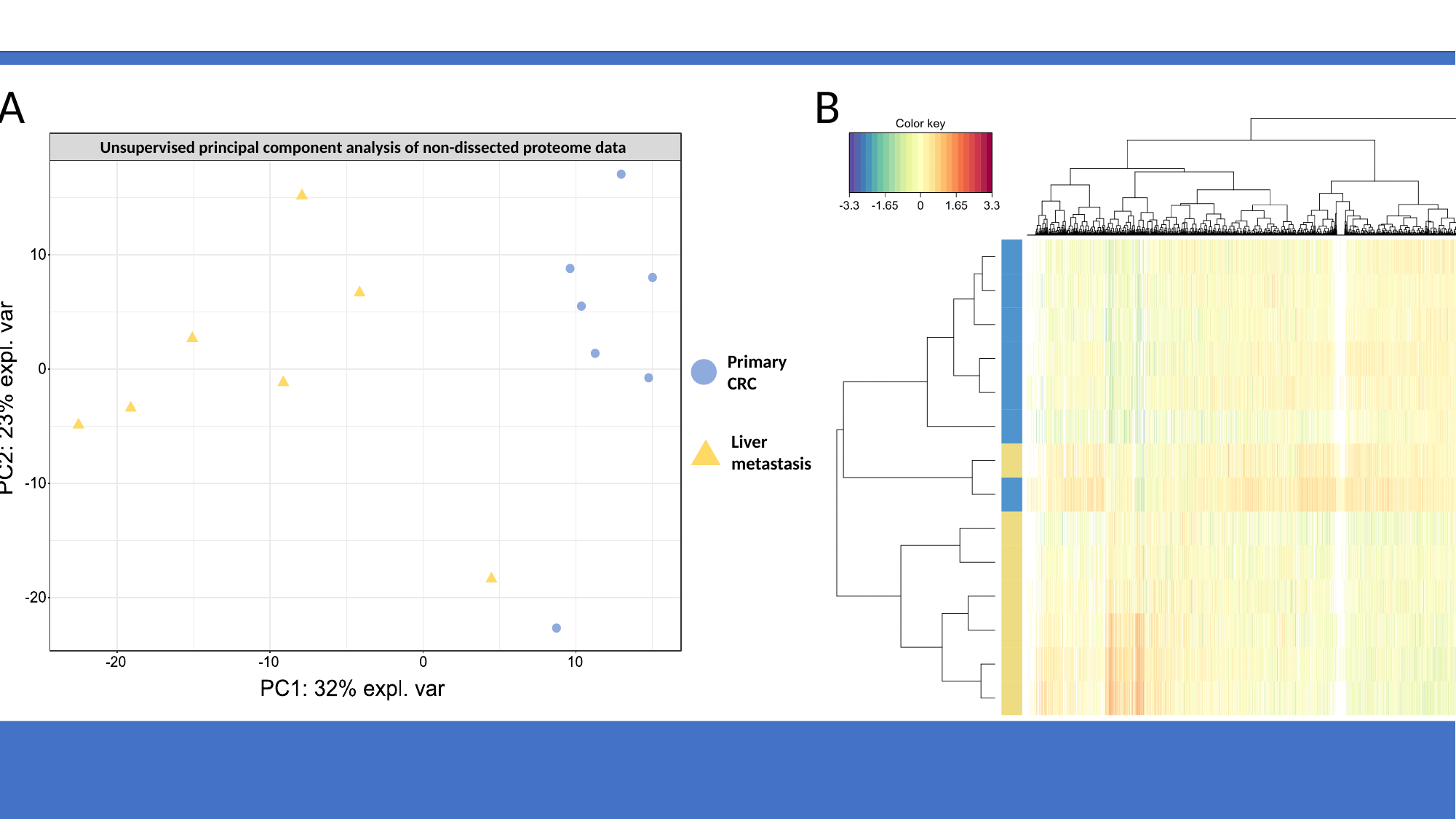

A
B
Unsupervised principal component analysis of non-dissected proteome data
Primary CRC
Liver metastasis
