## Supplementary Figure 4 for "Proteome biology of primary colorectal carcinoma and corresponding liver metastases"

### Slide 1
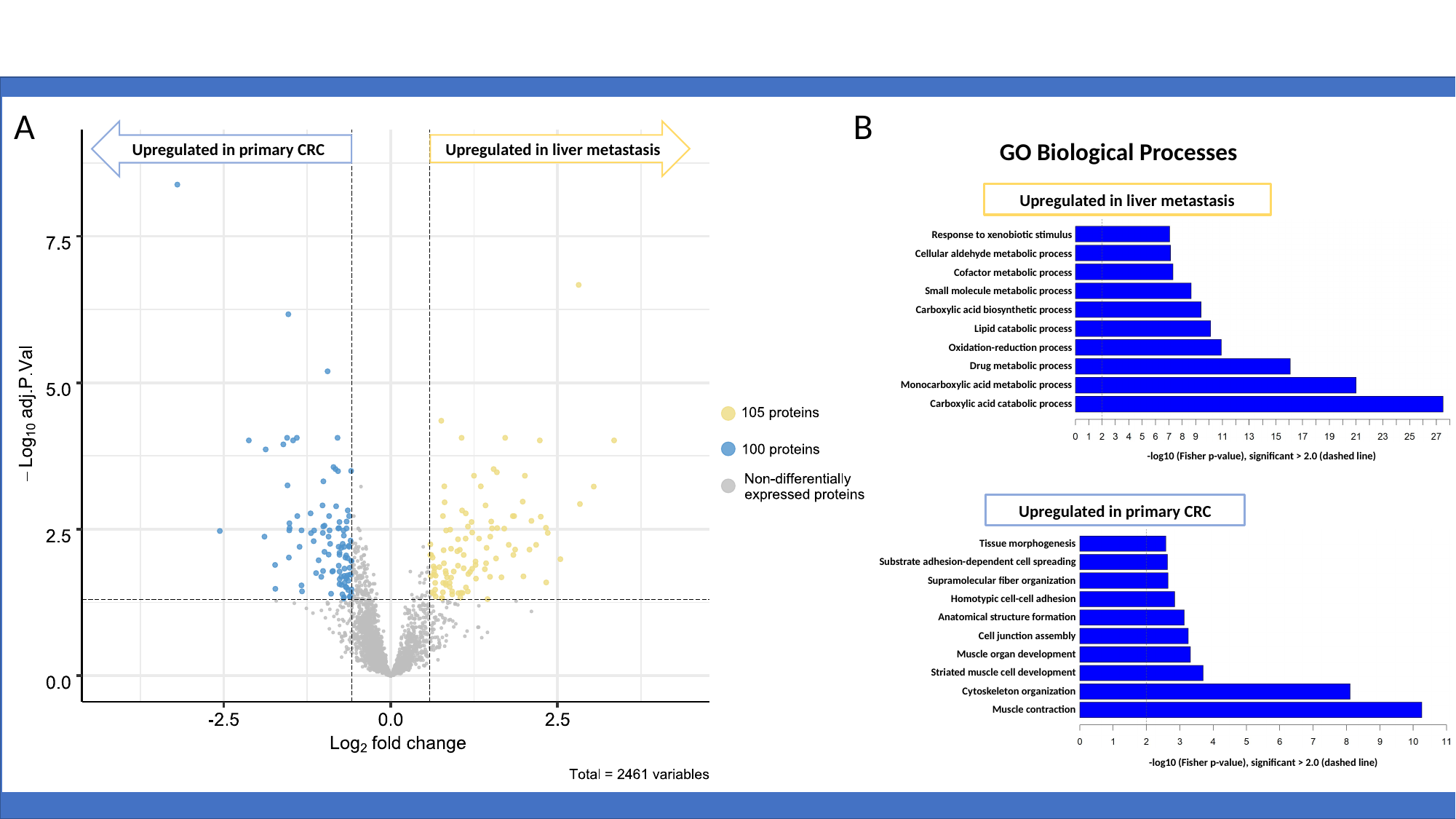

A
B
Upregulated in liver metastasis
Upregulated in primary CRC
GO Biological Processes
Upregulated in liver metastasis
Response to xenobiotic stimulus
Cellular aldehyde metabolic process
Cofactor metabolic process
Small molecule metabolic process
Carboxylic acid biosynthetic process
Lipid catabolic process
Oxidation-reduction process
Drug metabolic process
Monocarboxylic acid metabolic process
Carboxylic acid catabolic process
-log10 (Fisher p-value), significant > 2.0 (dashed line)
Upregulated in primary CRC
Tissue morphogenesis
Substrate adhesion-dependent cell spreading
Supramolecular fiber organization
Homotypic cell-cell adhesion
Anatomical structure formation
Cell junction assembly
Muscle organ development
Striated muscle cell development
Cytoskeleton organization
Muscle contraction
-log10 (Fisher p-value), significant > 2.0 (dashed line)
