## Supplementary Figure 5 for "Proteome biology of primary colorectal carcinoma and corresponding liver metastases"

### Slide 1
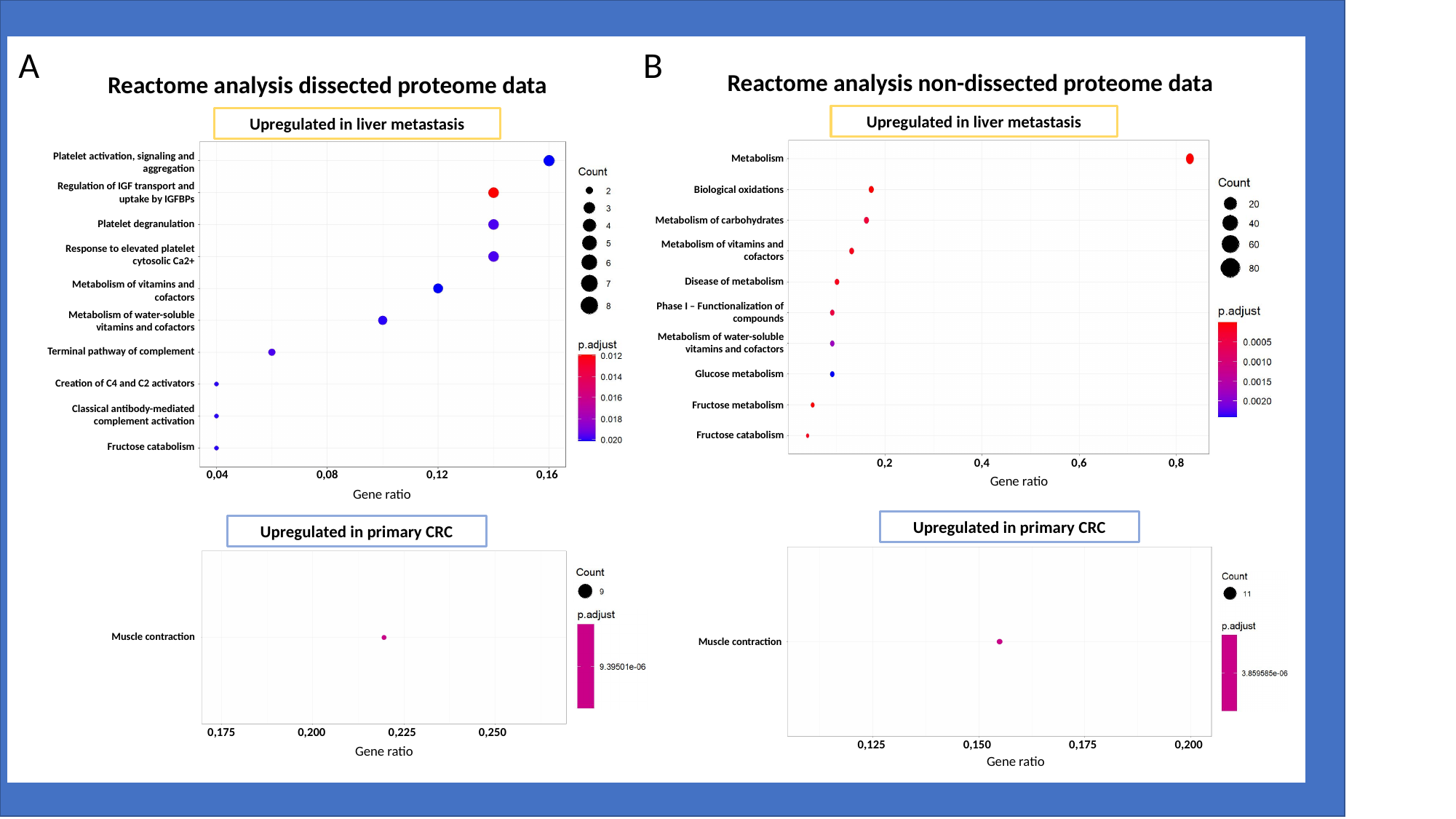

A
B
Reactome analysis non-dissected proteome data
Upregulated in liver metastasis
Metabolism
Biological oxidations
Metabolism of carbohydrates
Metabolism of vitamins and cofactors
Disease of metabolism
Phase I – Functionalization of compounds
Metabolism of water-soluble vitamins and cofactors
Glucose metabolism
Fructose metabolism
Fructose catabolism
0,2
0,4
0,6
0,8
Gene ratio
Upregulated in primary CRC
Muscle contraction
0,125
0,150
0,175
0,200
Gene ratio
Reactome analysis dissected proteome data
Upregulated in liver metastasis
0,04
0,08
0,12
0,16
Gene ratio
Platelet activation, signaling and aggregation
Regulation of IGF transport and uptake by IGFBPs
Platelet degranulation
Response to elevated platelet cytosolic Ca2+
Metabolism of vitamins and cofactors
Metabolism of water-soluble vitamins and cofactors
Terminal pathway of complement
Creation of C4 and C2 activators
Classical antibody-mediated complement activation
Fructose catabolism
Upregulated in primary CRC
Muscle contraction
0,175
0,200
0,225
0,250
Gene ratio
